# Interactive regulation between aliphatic hydroxylation and aromatic hydroxylation of thaxtomin D in TxtC: a theoretical investigation

**DOI:** 10.1101/2020.12.18.423417

**Authors:** Chang Yuan, Qingwen Ouyang, Xixi Wang, Xichen Li, Hongwei Tan, Guangju Chen

**Author notes:** **Corresponding Authors**: Hongwei Tan, Guangju Chen,. **Present Addresses** Key Laboratories of Theoretical and Computational Photochemistry, Ministry of Education, Beijing Normal University, Beijing, China.

## Abstract

TxtC is an unusual bifunctional cytochrome P450 that is able to perform sequential aliphatic and aromatic hydroxylation of the diketopiperazine substrate thaxtomin D in two remote sites to produce thaxtomin A. Though the X-ray structure of TxtC complexed with thaxtomin D revealed a binding mode for its aromatic hydroxylation, the preferential hydroxylation site is aliphatic C_14_. It is thus intriguing to unravel how TxtC accomplishes such two-step catalytic hydroxylation on distinct aliphatic and aromatic carbons and why the aliphatic site is preferred in the hydroxylation step. In this work, by employing molecular docking and molecular dynamics (MD) simulation, we revealed that thaxtomin D could adopt two different conformations in the TxtC active site, which were equal in energy with either the aromatic C-H or aliphatic C_14_-H laying towards the active Cpd I oxyferryl moiety. Further ONIOM calculations indicated that the energy barrier for the rate-limiting hydroxylation step on the aliphatic C_14_ site was 8.9 kcal/mol more favorable than that on the aromatic C_20_ site. The hydroxyl group on the monohydroxylated intermediate thaxtomin B C_14_ site formed hydrogen bonds with Ser280 and Thr385, which induced the _L_-Phe moiety to rotate around the C_β_−C_γ_ bond of the 4-nitrotryptophan moiety. Thus, it adopted an energy favorable conformation with aromatic C_20_ adjacent to the oxyferryl moiety. In addition, the hydroxyl group induced solvent water molecules to enter the active site, which propelled thaxtomin B towards the heme plane and resulted in heme distortion. Based on this geometrical layout, the rate-limiting aromatic hydroxylation energy barrier decreased to 15.4 kcal/mol, which was comparable to that of the thaxtomin D aliphatic hydroxylation process. Our calculations indicated that heme distortion lowered the energy level of the lowest Cpd I α-vacant orbital, which promoted electron transfer in the rate-limiting thaxtomin B aromatic hydroxylation step in TxtC.

## INTRODUCTION

Thaxtomins are a family of phytotoxic diketopiperazines and causative agents of multiple plant pathogenic Streptomyces strains that inhibit cellulose biosynthesis in expanding plant tissues.^1,2^ Thaxtomins are often used as herbicides in crop protection due to their virulence.^3,4^ However, the synthetic routes to thaxtomins suffer from a lack of stereocontrol and low overall yield.^5,6^ Recently, it was reported that TxtC is able to effectively produce thaxtomin A, an herbicide approved by the U.S. Environmental Protection Agency (EPA),^3,4,7^ from thaxtomin D. TxtC is a member of the cytochrome P450 family, which consists of versatile monooxygenases that perform a broad array of biochemical transformations, such as hydroxylation of alkanes, oxidation of heteroatoms or aromatics, and dealkylation reactions.^8–12^ In these catalytic reactions, the active high-valent iron-oxo porphyrin π-cation radical (Cpd I, as shown in Scheme S1) species serves as the oxygen source. The conversion from thaxtomin D to thaxtomin A in TxtC is a two-step process. First, Cpd I inserts an oxygen atom into the thaxtomin D aliphatic C_14_-H bond to generate thaxtomin B, which is followed by the second hydroxylation step on the thaxtomin B phenyl group meta-position to produce thaxtomin A^3,4,7^ (see Scheme 1A). It is worth noting that aliphatic and aromatic C-H bonds usually have energy differences as high as 10~20 kcal/mol in the bond dissociation energies (BDEs), which means that TxtC is able to use distinct strategies to accomplish sequential aliphatic and aromatic hydroxylation on thaxtomin D. The cytochrome P450 catalytic hydroxylation mechanism has been extensively investigated by both experimental and theoretical studies. As shown in Scheme 1B, aliphatic hydroxylation by cytochrome P450 follows a common mechanism that involves initial hydrogen atom abstraction (HAT) by Cpd I followed by hydroxyl rebounding.^13–18^ Aromatic hydroxylation by cytochrome P450 is much more complicated.^16–20^ Scheme 1C summarizes several mechanistic hypotheses. All the mechanisms leading to hydroxylation on the aromatic ring are channeled through a tetrahedral σ-complex intermediate (INT2 in Scheme 1C) formed by oxyferryl from Cpd I added to the substrate aromatic carbon. The subsequent reaction process may follow different pathways. One of the proposed pathways (Pathway-A, Scheme 1C) goes through an epoxide intermediate, which then converts to a hydroxylation product by cleaving the oxygen and the ortho-carbon bond accompanied by a 1,2-hydrogen shift (NIH shift)^21^. However, deuterium isotope experiments indicated that aromatic hydroxylation by cytochrome P450 is unlikely to follow such an epoxide formation mechanism.^22^ An alternative aromatic hydroxylation mechanism via the tetrahedral σ-complex intermediate was proposed by Shaik et al.^20^ (Pathway-B, Scheme 1C), in which the from the substrate tetrahedral carbon to the oxyferryl moiety oxygen atom through one of the pyrrole N in the porphyrin ring. Proton migration from the tetrahedral carbon to the oxygen atom could also be accomplished through the NIH shifting mode, as shown by pathway-C in Scheme 1C in which the proton shuttle is mediated by the ortho-carbon atom of the phenyl ring. Both aliphatic and aromatic hydroxylation are well within the scope cytochrome P450 activities, but how TxtC integrates these two activities in a single active center is not yet understood. Further effort is therefore needed to unravel the bifunctional TxtC hydroxylation mechanism.

**Scheme 1.**
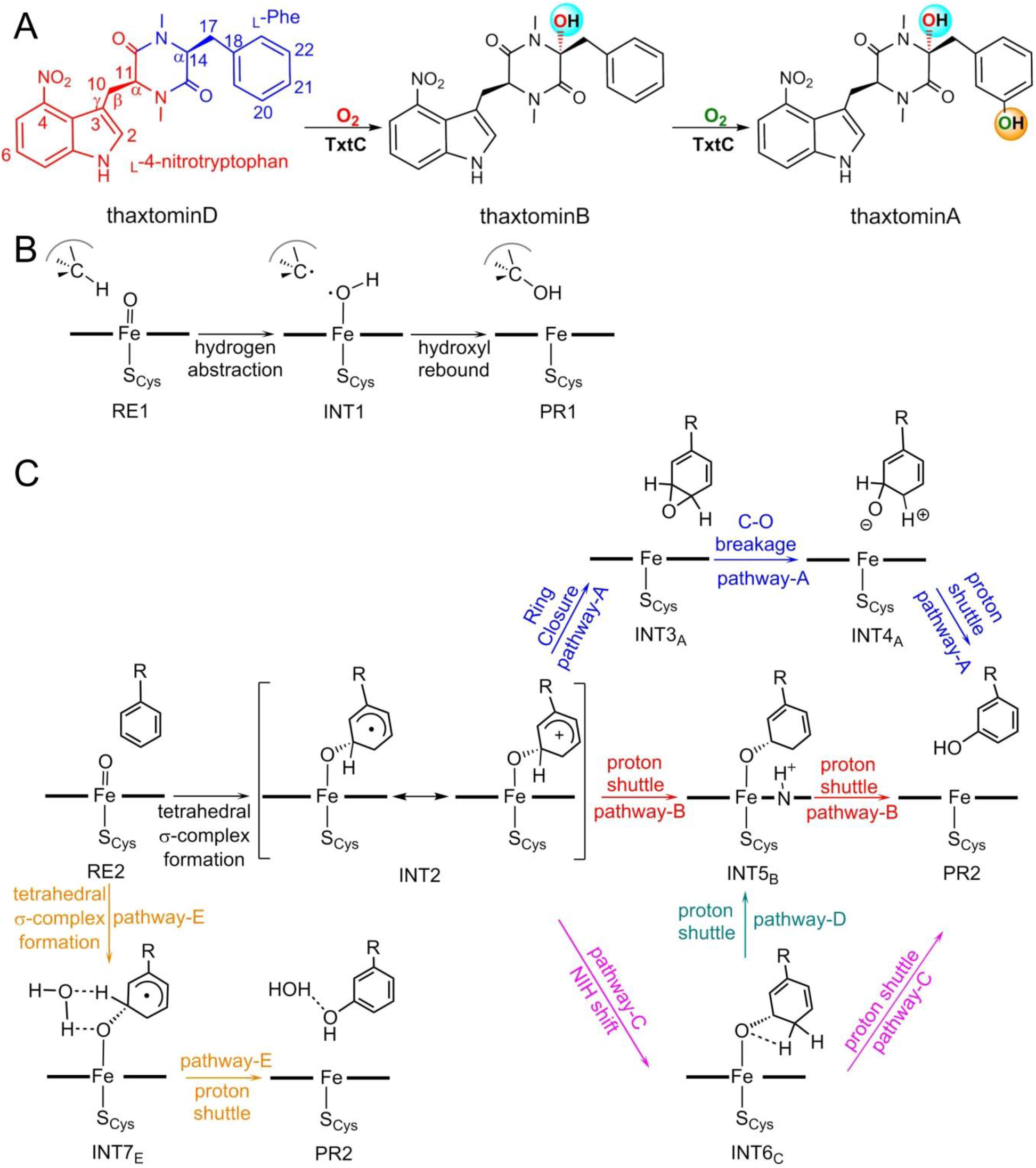
(A) Thaxtomin A biosynthetic route, starting with the thaxtomin D which is assembled from _L_-Phe and _L_-4-nitrotryptophan. Alternative mechanistic hypotheses for (B) aliphatic hydroxylation and (C) aromatic hydroxylation by cytochrome P450.

Recently, Challis et al. determined the X-ray crystal structures of TxtC complexed with thaxtomin D (PDB code: 6f0b) and thaxtomin B (PDB code: 6f0c).^7^ As shown in Figure 1A and 1B, heme functions as the prosthetic group in these two complexes, of which the iron cation connects to the protein backbone via the distal Cys ligand thiolate linkage (Cys344). It is interesting to note that in complex structures, thaxtomin D adopts a conformation similar to thaxtomin B. Both substrate molecules place the phenyl group in the vicinity of the oxyferryl moiety, though the preferential thaxtomin D hydroxylation site is its aliphatic C_14_ atom. Therefore, thaxtomin D should have an alternative conformation in TxtC’s active site by placing C_14_ next to the catalytic iron center. Docking calculations conducted by Challis et al. supported the speculation that the TxtC active site can accommodate two distinct thaxtomin D binding scenarios, in which either the thaxtomin D phenyl group or aliphatic C_14_ lies over the heme plane^7^. Given that TxtC clearly displays sequential hydroxylation of aliphatic C_14_ prior to the aromatic C_20_ on thaxtomin D, an intriguing question is how TxtC realizes such regioselectivity. Several previous studies have investigated the cytochrome P450 regioselective oxidation reaction. For the 1-methyl-4-phenyl-1,2,3,6-tetrahydropyridine (MPTP) detoxification process by cytochrome P450 (mainly CYP2D6), theoretical calculations identified that N-demethylation is thermodynamically more favorable than the aromatic hydroxylation process.^23^ Experimental results from synthesized biomimetic iron–oxo porphyrin oxidants indicated that the iron proximal axial ligand is an important factor in oxidation regioselectivity. With acetonitrile as the proximal ligand, [Fe^IV^=O(Por^+^·)NCCH_3_] showed aromatic preference in the ethylbenzene oxidation reaction, while [Fe^IV^=O(Por^+^·)Cl^−^] predominantly produced 1-phenylethanol as the oxidation product.^24^ A follow-up DFT study revealed that the anionic axial ligand chloride weakens the oxidant electrophilicity so that the aromatic hydroxylation process, which depends on electron transfer from the substrate to the oxidant, is significantly suppressed.^25^ In addition, Shaik et al. indicated that the hydrogen bonds formed around the axial thiolate ligand act as an external electric field along the S-Fe-O axis and change the electronic Cpd I conformation, thus dictating the regioselectivity between epoxidation and allylic hydroxylation of propene by Cpd I.^26^ Generally, as an active oxidant, Cpd I acts as a “chemical chameleon” owing to its maneuverable electronic structure in different protein environments. However, it is not clear whether and how the TxtC structural and electronic factors determine its regioselectivity between thaxtomin D aliphatic and aromatic hydroxylation. In fact, TxtC performs aromatic hydroxylation on the hydroxylated intermediate thaxtomin B instead of thaxtomin D, and whether the hydroxyl moiety on the thaxtomin B C_14_ site influences the aromatic hydroxylation reaction process remains unknown. Kinetic experiments determined that thaxtomin B aromatic hydroxylation efficiency is on the same order as that of thaxtomin D aliphatic hydroxylation in TxtC,^3^ which implies that TxtC uses an unknown mechanism to regulate the two-step hydroxylation reactions.

**Figure 1.**
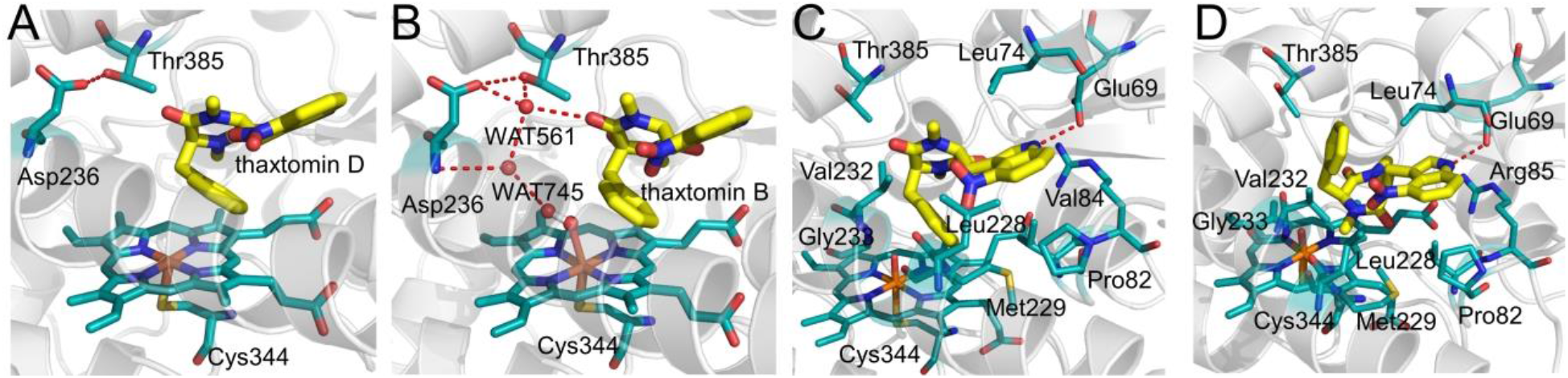
Active sites of TxtC combines with (A) thaxtomin D (PDB code: 6f0b) and (B) thaxtomin B (PDB code: 6f0c) in the X-ray crystal structures. MD simulated complexes of TxtC-thaxtomin D in (C) complex A and (D) complex B. The key residues in the active site are represented as sticks. Carbon, oxygen, sulfur, and nitrogen atoms are colored in cyan, red, orange, and blue, respectively. The carbon atoms in the substrate molecules are colored in yellow. The H-bonds are represented as red dashed lines.

In this study, molecular dynamics (MD) simulations and ONIOM calculations were employed to address the mechanistic details of sequential thaxtomin D aliphatic and aromatic hydroxylation in TxtC. Our computational results revealed that the initial thaxtomin D aliphatic C_14_–H activation proceeded via a rate-limiting HAT step with a hydrogen atom on C_14_ transferring to the Cpd I oxoferryl, which was followed by the hydroxyl moiety rebound to generate the C_14_ hydroxylated intermediate thaxtomin B. The hydroxyl group on the C_14_ site formed hydrogen bonds with Ser280 and Thr385, which induced thaxtomin B isomerization to place the aromatic carbon (C_20_) next to the oxyferryl moiety. This initiated the three-step thaxtomin B aromatic hydroxylation: (1) the rate-limiting tetrahedral σ-complex intermediate formation with electron transfer from thaxtomin B to Cpd I, (2) NIH shift, and (3) proton shuttling via porphyrin. Our ONIOM calculation and molecular dynamics simulation results showed that the thaxtomin B C_14_ hydroxyl group played a crucial role in its aromatic hydroxylation process by inducing bulk water molecules to enter the active pocket, which propelled thaxtomin B towards the heme plane and resulted in out-of-plane heme ring distortion. Such a conformational change promoted Cpd I electrophilicity in TxtC so that it was able to easily perform aromatic hydroxylation on thaxtomin B. This work revealed the critical role of the hydroxyl moiety on the thaxtomin B C_14_ site in steering its aromatic C-H hydroxylation and provided new insights into the interactive influence between the substrate and enzyme activity.

## MODELS AND METHODS

The TxtC-thaxtomin D complex X-ray structure showed that the substrate adopted an orientation with which Cpd I could not perform hydrogen atom extraction from the C_14_ site.^7^ To obtain a proper starting structure of thaxtomin D complexed in TxtC for aliphatic carbon hydroxylation, molecular docking was conducted based on the TxtC crystal structure (PDB code: 6f0b)^7^ by using the program AutoDock 4.2.^27^ When performing the docking calculations, the TxtC protein was set to be rigid, while for the substrate thaxtomin D, all the C-C and C-N single bonds were set to be rotatable to better fit in the pocket. A cubic grid with 108*90*108 grid points and a spacing of 0.59 Å was chosen to ensure complete TxtC coverage. The docking simulation was performed with 500 cycles using the Lamarckian genetic algorithm (LGA).^28^

Docking calculations offered two optimal TxtC-thaxtomin D complex structures in which thaxtomin D placed its phenyl group and aliphatic C_14_ next to the oxyferryl moiety (Figure S1). These two complexes of TxtC-thaxtomin D and TxtC-thaxtomin B (taken from X-ray structure with PDB code: 6f0c) were then submitted for further MD simulations. Hydrogen atoms were added to the complex structures by using the PROPKA program^29^ based on the experimental pH environment. The Amber ff14SB force field^30^ was used for the protein, while the substrate molecules thaxtomin D and thaxtomin B were parametrized using the ANTECHAMBER module.^31,32^ The heme oxyferryl complex (Cpd I) force field parameters were taken from the literature^33^. The three complexes were soaked in a TIP3P^34^ water box with a minimum distance of 10 Å to the protein boundary and neutralized with Na^+^ counterions. Each of the processed systems was then subjected to a three-step energy minimization process so the hydrogen atoms, solvent molecules, and the whole system could prevent possible clashes. After energy minimization, the systems were then heated to 300 K and equilibrated for 300 ps. Finally, MD simulations were performed in the NPT ensemble for 100 ns to obtain the product phase.

From each of the MD simulation trajectories, three independent and stable snapshots at 50, 80, and 100 ns were abstracted and used to build the models for mechanism investigation with the ONIOM approach.^35^ Since the TxtC protein was rather large, a truncated model that contained the residues within 10 Å of the active site was built for ONIOM calculations. The QM region of the model consisted of (1) the active oxidant Cpd I, (2) the thiolate cysteine residue (Cys344), which was the iron center proximal ligand, and (3) the substrate (thaxtomin D/thaxtomin B). In the TxtC-thaxtomin B model, two crystal water molecules (WAT561 and WAT745) residing in the active site and having hydrogen bond interactions with Asp236 and Thr385 were also included. The QM-MM layer boundary was captured with hydrogen atoms. The atoms in the MM region, except those within 6 Å of the QM zone, were frozen during optimization. The TAO toolkit^36^ was employed in preparing the ONIOM calculations.

All geometry optimizations, including transition structure searches, were performed by using the ONIOM protocol, in which the DFT hybrid functional B3LYP^37^ with def2-SVP^38^ basis set was employed for the QM atoms, while the AMBER ff98 force field^39^ embedded in the Gaussian 09 program^40^ was used for the MM part. Intrinsic reaction coordinate (IRC) and harmonic vibrational frequency calculations were conducted at the same theoretical optimization level to confirm correct stationary point structures locations. Zero-point energy (ZPE) and entropy corrections were obtained based on the frequency calculations. Van der Waals effect was assessed by using Grimme’s D3 method.^41^ Energy corrections with higher accuracy were then evaluated with the ONIOM (electronic embedding) protocol by using B3LYP/def2-TZVP^42^ for the QM atoms. All calculations were performed with the Gaussian09 program package.^40^

## RESULTS AND DISCUSSION

### The target systems

TxtC performs sequential hydroxylation on the thaxtomin D aliphatic C_14_ and thaxtomin B aromatic C_20_, while the X-ray structure of thaxtomin D complexed in TxtC revealed a conformation with its aromatic C_20_ was next to the heme iron center. Thus, thaxtomin D could adopt an alternative conformation in the TxtC active site to fulfil its aliphatic C_14_ carbon hydroxylation. Molecular docking was thus performed to obtain the possible thaxtomin D binding conformation for its C_14_ hydroxylation. The docking results showed that the TxtC-thaxtomin D complex indeed contained a mixture of two thaxtomin D conformations. In docked complex A (see Figure S1A), thaxtomin D adopted a conformation similar to that in the X-ray structure (PDB code: 6f0b),^7^ while in the second docked complex (complex B, Figure S1B), thaxtomin D lies at its aliphatic hydroxylation site C_14_ towards the oxyferryl moiety. These docking results were consistent with those from Challis et al.^7^ Further 100 ns MD simulations on the two complexes showed converged root-mean-squared deviations (RMSDs, as shown in Figure S2), which confirmed that they were both stable binding states. Moreover, MM-PBSA calculations identified that thaxtomin D had a binding energy similar to TxtC in the two complexes (−45.3 kcal/mol vs. −44.7 kcal/mol), which further confirmed that thaxtomin D could adopt two energetically equal conformations in the TxtC active site. As shown in Figure 1C, in complex A, the thaxtomin D phenyl group was placed over the oxyferryl moiety from the γ-side of the heme. The substrate _L_-4-nitroindole moiety, which formed a hydrogen bond with the residue Glu69, was situated in the hydrophobic pocket formed by Leu74, Pro82, Val84, Arg85, Leu228, and Met229. The _L_-Phe part of the substrate had hydrophobic contact with residue Thr385. Compared to that in complex A, thaxtomin D twisted around the C_β_−C_γ_ bond of the 4-nitrotryptophan moiety by ~123° in complex B so that its phenyl group turned to point towards the upper active pocket (Figure 1D). Such a conformation allowed thaxtomin D C_14_ to approach the TxtC oxyferryl moiety at a close distance. In complex B, the substrate _L_-4-nitrotryptophan group remained in the hydrophobic pocket formed by Leu74, Pro82, Val84, Arg85, Leu228, and Met229 similar to that in the X-ray structure. Such a geometrical layout of complex B ensured reasonable orientation for the observed thaxtomin D aliphatic C_14_ hydroxylation regiochemistry. Complex B was then employed to investigate the thaxtomin D aliphatic hydroxylation mechanism in TxtC.

### Thaxtomin D aliphatic hydroxylation by TxtC

Considering that the active species Cpd I was in the form of triradicaloids^17,18^, the sequential tandem thaxtomin D hydroxylation processes were investigated by two-state reactivity involving doublet (S=1/2) and quartet (S=3/2) spin states with three different snapshots from MD trajectories as the reactants. As displayed in Figure 2, our calculations showed that the thaxtomin D aliphatic C_14_ hydroxylation reaction proceeded on the doublet state potential energy surface through a rate-limiting hydrogen abstraction step with a 14.6 kcal/mol energy barrier (By using the other two snapshots as the reactant initial structures, 14.5 kcal/mol and 14.0 kcal/mol energy barriers were obtained), which was followed by a hydroxyl radical rebound step. The following discussion of the thaxtomin D aliphatic hydroxylation reaction in TxtC is focused on the energetically dominant doublet state.

**Figure 2.**
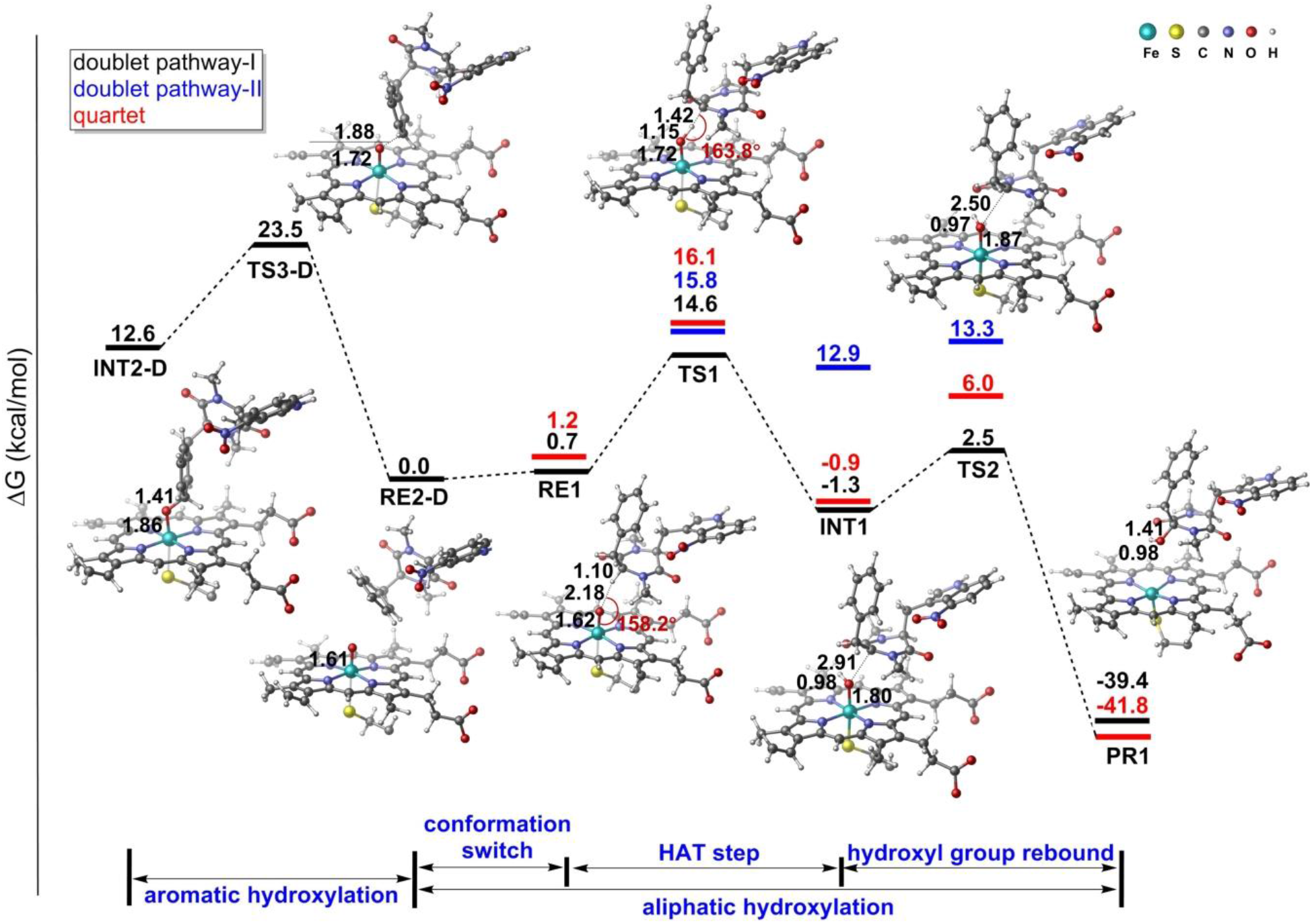
Energy profile of the TxtC-catalyzed thaxtomin D aliphatic and aromatic hydroxylation processes. The key structures of the stationary point in the doublet state are shown. Bond lengths are given in Å.

The thaxtomin D aliphatic hydroxylation process started from reactant ^2^RE1, in which the thaxtomin D C_14_ hydrogen atom pointed towards the Cpd I oxyferryl moiety with a H···O distance of 2.18 Å and a ∠C−H−O angle of 158.2° (Figure 2). Table 1 displays the spin distribution of the important reaction species during the C_14_ hydroxylation process. Based on the spin population, in ^2^RE1 two unpaired α electrons resided between the oxyferryl moiety, which antiferromagnetically coupled with one unpaired β electron residing on the proximal Cys344 ligand. In common Cpd I, the porphyrin ring usually presents as a cationic radical, while in ^2^RE1, the spin population summed on the porphyrin ring is less than −0.30. Such a spin population indicated that the anionic Cys344 ligand was partially oxidized by the porphyrin moiety in ^2^RE1.

**Table 1.** The spin population of doublet stationary points in thaxtomin D aliphatic hydroxylation in TxtC. In this table, O, Por, thxD, and H_C14_ represent the oxo moiety of Cpd I, porphyrin, thaxtomin D, and hydrogen attached C_14_, respectively.

| Compound | Fe | O | Cys344 | Por | thxD | H<sub>C14</sub> | C<sub>14</sub> |
| --- | --- | --- | --- | --- | --- | --- | --- |
| <sup>2</sup>RE2 | 1.33 | 0.76 | -0.79 | -0.30 | 0.01 | 0.00 | 0.00 |
| <sup>2</sup>TS1 | 2.07 | 0.13 | -0.39 | -0.20 | -0.61 | 0.02 | -0.39 |
| <sup>2</sup>INT1 | 1.84 | 0.21 | 0.07 | -0.16 | -0.96 | 0.01 | -0.62 |
| <sup>2</sup>TS2 | 1.62 | 0.21 | 0.01 | -0.14 | -0.71 | 0.00 | -0.45 |
| <sup>2</sup>PR1 | 1.23 | 0.00 | -0.10 | -0.14 | 0.00 | 0.00 | 0.00 |

The thaxtomin D conformation in ^2^RE1 allowed oxyferryl to selectively abstract hydrogen atoms from C_14_ via transition state ^2^TS1. In the optimized ^2^TS1, C_14_-H-O tended to have a collinear arrangement with C_14_-H bond lengths and O···H distances of 1.42 Å and 1.15 Å, respectively, showing a typical feature of the C-H bond activation transition state by Fe^IV^=O species.^43–45^ From ^2^RE1 to ^2^TS1, along with the C-H bond stretching, an electron is partially transferred from the substrate to the (Por^+^-Cys^−^)**·** moiety, evidenced by the calculated NPA charge on the substrate increasing to +0.41 in ^2^TS1 (Table S1). The spin population on the substrate decreased to −0.61, while that on the (Por^+^-Cys^−^)**·** moiety increased to −0.59 from −1.09 in ^2^RE1, showing that an electron in the α spin was being transferred from the substrate to the Por^+^-Cys^−^ moiety (pathway-I), as shown in Scheme S2. It therefore leads us to conclude that this reaction step proceeded via a hydrogen atom transfer (HAT) mechanism. This HAT process was the rate-limiting step of the whole C_14_ aliphatic hydroxylation reaction with a 14.6 kcal/mol energy barrier, which generated transient intermediate ^2^INT1. ^2^INT1 was a typical ferryl-hydroxyl species, in which the Fe-O bond length was measured as 1.80 Å. As shown in Table 1, the calculated spin populations on the iron and O atoms were 1.84 and 0.21, respectively, in ^2^INT1, corresponding to an intermediate spin state Fe^IV^-O^2−^. C_14_ presented as an alkyl radical with a spin population of −0.62. The following hydroxyl radical rebound step only needed to cross over a small local barrier of 3.8 kcal/mol and generated the TxtC-thaxtomin B complex (P1).

Following a similar reaction mechanism, the C_14_ hydroxylation reaction could also proceed through another possible mode with β spin electrons, but not α spin electron transferred from the substrate to Cpd I during the HAT process (pathway-II, see Scheme S2). As shown in Figure 2, pathway-II was less energetically favorable than pathway-I. Spin population analysis indicated that this β electron was transferred to the Fe^IV^=O moiety during HAT. In pathway-I, the α electron from the substrate went to the (Por^+^-Cys^−^)**·** part, which quenched the (Por^+^-Cys^−^)**·** radical, thus resulting in a favorable electron transfer process in the HAT reaction.

### Thaxtomin B conformation conversion in the TxtC active site

After the C_14_ hydroxylation reaction, to continue hydroxylation on the aromatic C_20_ site, the intermediate thaxtomin B needed to change its conformation to place its aromatic C_20_ site next to the oxyferryl moiety. However, whether thaxtomin B accomplished such a conformational switch by releasing-rebinding or directly flipping over inside the active pocket remains unclear. To investigate how thaxtomin B realized such a conformational switch, MD simulation was performed on the TxtC-thaxtomin B complex in which thaxtomin B adopted the conformation with its hydroxylated C_14_ in close proximity to the heme iron atom, which resembled the product state after C_14_ hydroxylation. The TxtC-thaxtomin B complex was then soaked in a TIP3P water box and submitted for 100 ns MD simulation. Interestingly, an obvious thaxtomin B interconversion from its initial conformation to the one with its phenyl groups laying over the iron center was observed in the MD simulation. The thaxtomin B conformational switch was characterized by the rotation of the _L_-Phe group around the C_β_−C_γ_ bond of the _L_-4-nitrotryptophan moiety, which changed significantly during the simulation so that the phenyl group flipped to lie over the heme plane (see Figure 3A). Correspondingly, the complex system energy decreased throughout the simulation (Figure S3). However, such conformational interconversion was not observed in the TxtC-thaxtomin D complex (complex B) 100 ns MD simulation (Figure 3A). This result therefore indicated that the C_14_ hydroxyl group was the driving force for the substrate thaxtomin B conformational switch.

**Figure 3.**
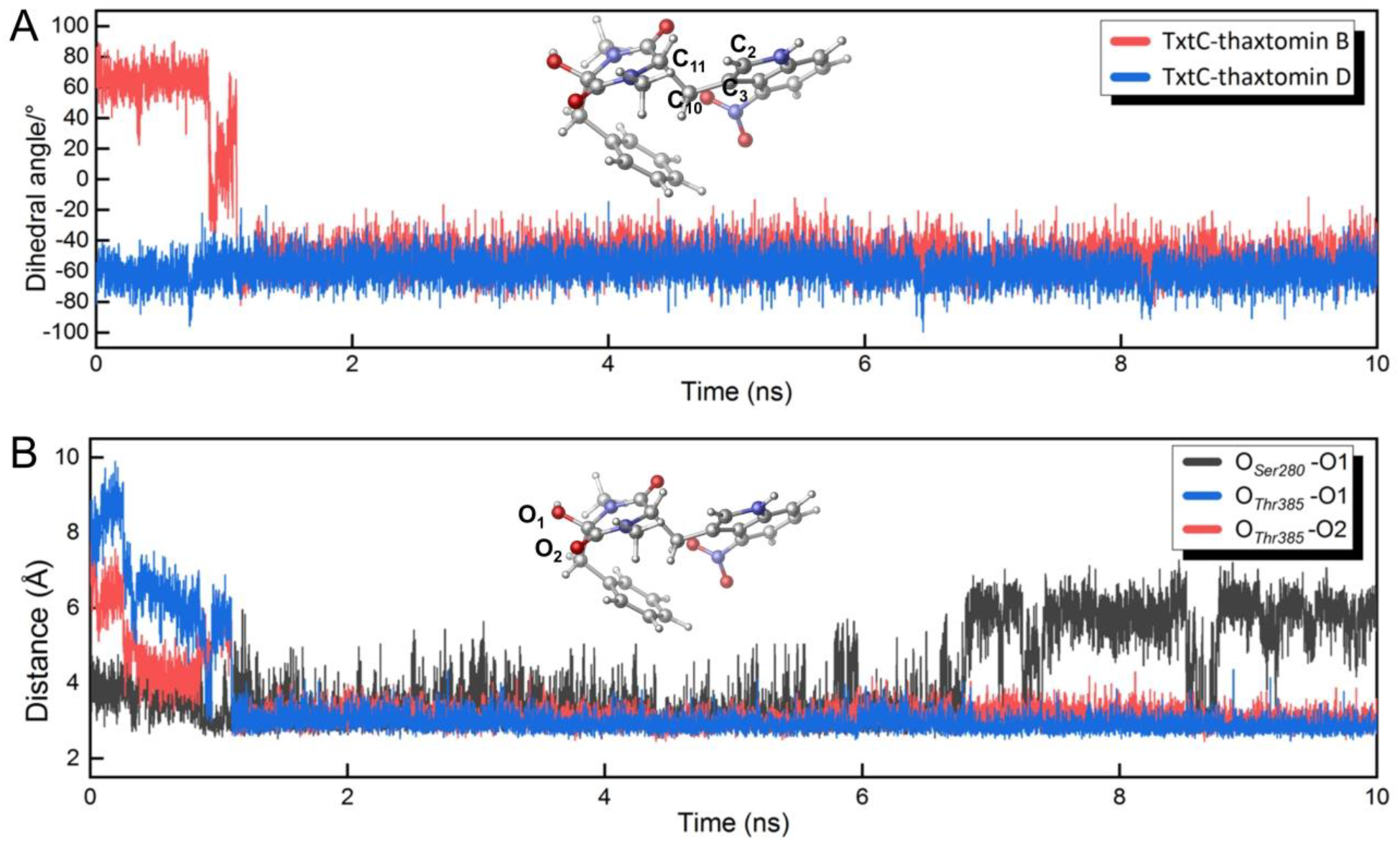
(A) Time-dependences of the thaxtomin B and thaxtomin D C_2_-C_3_-C_10_-C_11_ dihedral angle variations. (B) The distance variations of the hydrogen bonds between the thaxtomin B and Ser280 and Thr385 in 10 ns MD simulations (in complex B).

As shown in Figure 3B and Figure 4A, in the initial 10 ns MD simulation, Ser280 displayed a significant displacement. In the initial complex, Ser280 did not interact with the substrate but formed a hydrogen bond with Gly366 through the β-hydroxyl (O_Ser280_-O_Gly336_ = 2.96 Å). At 10 ps, Ser280 abolished its hydrogen bond interaction with Gly366 and formed a new hydrogen bond with the carbonyl group on thaxtomin B. As shown in Figure 4A, this hydrogen bonding interaction pulled thaxtomin B to deviate from its initial position and further resulted in C_14_-OH hydrogen bonding interactions with Ser280 (O_Ser280_-O_1_ distance = 3.25 Å) (11 ps, Figure 4A). At 892 ps, Ser280 exclusively interacted with C_14_-OH. Through hydrogen bonding with Ser280, the C_14_-OH group approached Thr385 at 903 ps. C_14_-OH turned to a hydrogen bond with Thr385 exclusively at 1102 ps. Such a relaying H-bonding interaction between C_14_-OH and the residues around it led to the entire thaxtomin B molecule being lifted away from the heme plane with a space between them. This allowed the _L_-Phe moiety to rotate around the 4-nitrotryptophan C_β_−C_γ_ bond and finally switch to the conformation with the phenyl group laying over the heme plane. Such a thaxtomin B conformation was similar to that in the TxtC-thaxtomin B X-ray structure. With the phenyl group flipping over, Ser280 departed from the C_14_-OH group, which then gradually returned back to its initial position and restored its hydrogen bonding interaction with Gly336. As a cross-validation, we also performed a 100 ns MD simulation on the TxtC-thaxtomin B complex with its initial conformation taken from the X-ray structure (PDB code: 6f0c) in which the thaxtomin B phenyl group lay on the heme plane. Thaxtomin B conformational interconversion was not observed during the simulation (Figure 4B). It therefore led us to conclude that (1) thaxtomin D had two equivalent binding conformations in TxtC’s active site; (2) hydroxylation at the C_14_ site induced thaxtomin B conversion to reorient its phenyl group towards the oxyferryl moiety; and (3) the conformation conversion was irreversible, which enabled the second hydroxylation on the aromatic C_20_ site to proceed on a stable thaxtomin B binding state.

**Figure 4.**
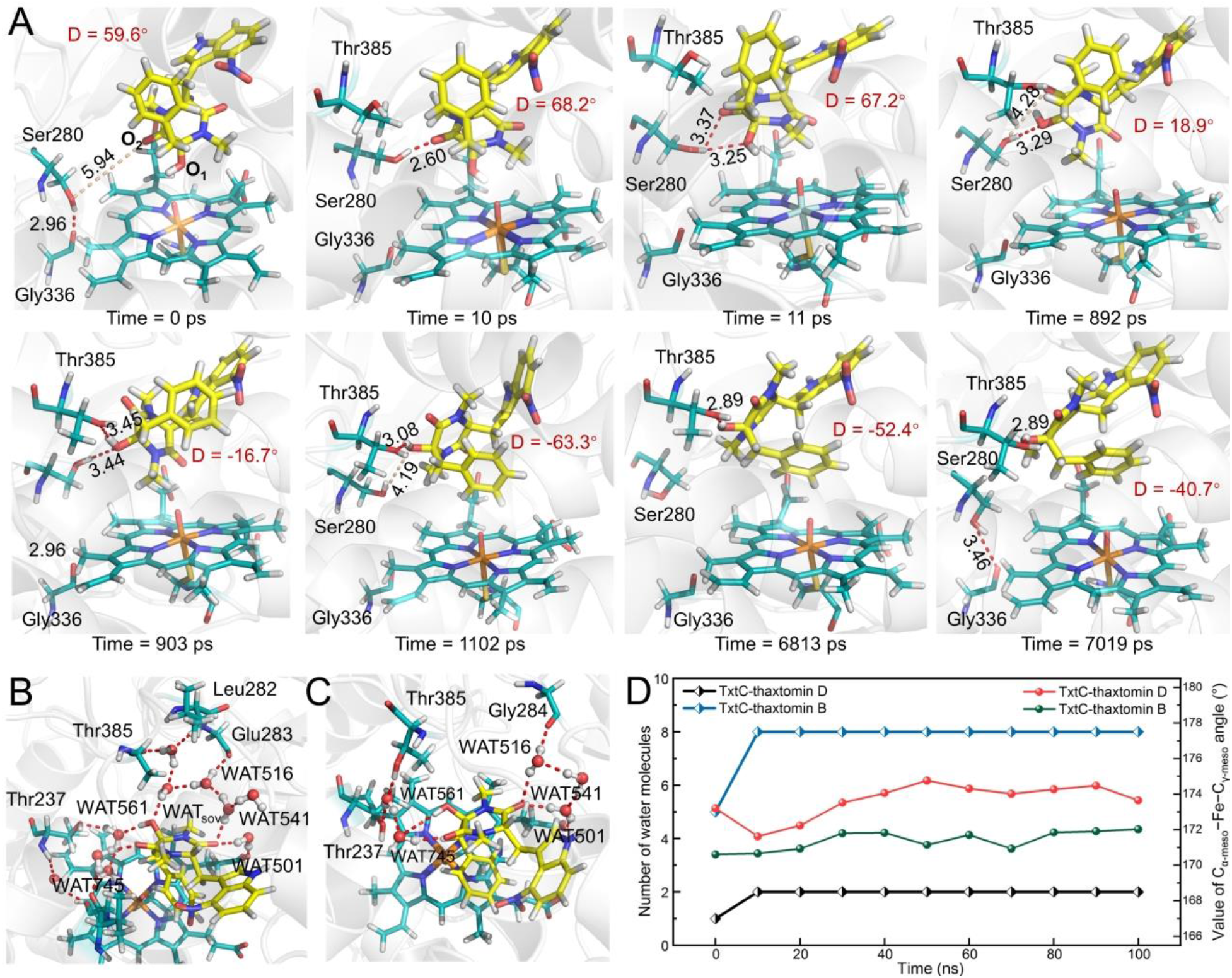
(A) The representative structures in the conformational switch of thaxtomin B from complex B to complex A along the MD simulation. TxtC-thaxtomin B in its X-ray structure after (B) 100 ns MD simulation and (C) the starting structures. (D) The quantity of water variation in the binding cavities of thaxtomin D and thaxtomin B were counted with interval setting as 10 ns. The key residues in the active site are represented as sticks. Carbon, oxygen, sulfur, and nitrogen atoms are colored in cyan, red, orange, and blue, respectively. The carbon atoms in the substrate molecules are colored in yellow. The H-bonds are represented as red dashed lines.

The hydroxyl group on the C_14_ site not only caused thaxtomin B conformational conversion, but also reshaped the protein environment around it. By comparing the MD simulated complexes of TxtC-thaxtomin D (complex A, from molecular docking) and TxtC-thaxtomin B (from X-ray structure, PDB code: 6f0c), we found subtle differences between them, although the substrate molecules were oriented similarly in the two complexes. As shown in Figure 4B, in the TxtC-thaxtomin B complex, the polar hydroxyl on C_14_ changed its surrounding environment and induced water molecules to enter the active site. An intuitive comparison between the two complexes is given in Figure 4D, in which we plotted the number of water molecules in the TxtC binding cavities during 100 ns MD simulations. Figure 4B and 4C depict two snapshots (100 ns and 0 ns) of TxtC-thaxtomin B complexes during MD simulation that represent the product and initial structures, respectively. There were two separate hydrogen bond networks formed on the interface between thaxtomin B and TxtC in the initial structure. The one on the left side of Figure 4C consisted of two water molecules (WAT561 and 745), residues Thr237 and Thr385, and the thaxtomin B carbonyl group. The 4-nitrotryptophan moiety carbonyl group, three water molecules (WAT501, 516 and 541), residue Gly284, and one of the carboxylate side chains on the heme formed the other hydrogen bond network in the active site, as shown on the right side of Figure 4C. After 100 ns MD simulation, Figure 4B shows that these two hydrogen bond networks merged because several bulk water molecules joined them around the C_14_ hydroxyl group. However, the water molecules present in the TxtC-thaxtomin D complex active site during MD simulation were far fewer than those in the TxtC-thaxtomin B complex. As one can see from Figure S4, for the TxtC-thaxtomin B complex, at the beginning of the MD simulation there was a water molecule (WAT421) in the active site that bridged the thaxtomin B 4-nitrotryptophan carbonyl with a heme carboxylate side group. In the 100 ns MD simulation, on average, only one water molecule entered into the active site from the bulk solvent. Such an observation clearly reflected the hydrophobicity of the interface between thaxtomin D and TxtC, which, nevertheless, changed its hydrophilicity with the hydroxyl group being induced on the thaxtomin B C_14_. Though the water molecules induced into the active site by C_14_-OH did not cause any significant protein tertiary structure change, they indeed resulted in profound conformation change on the heme plane. These water molecules crowded the active site and forced the thaxtomin B molecule to move towards the heme ring, of which the phenyl group was pushed to stack on the heme γ-edge. The steric interaction between the thaxtomin B phenyl group and the heme ring caused the heme γ-edge to bend towards the proximal ligand Cys344 and consequently led to obvious heme out-of-plane distortion. The angle C_α-meso_-Fe-C_γ-meso_ could be used to measure the heme plane distortion degree. During the 100 ns simulation, the average TxtC-thaxtomin B C_α-meso_-Fe-C_γ-meso_ angle was 171.5°, which deviated from 180.0° of a planar heme by 8.5°. Thaxtomin D showed less deformation in TxtC with the corresponding C_α-meso_-Fe-C_γ-meso_ angle of 173.8° during MD simulation. Heme distortion could profoundly change the enzyme reactivity and even the catalytic mechanism, which has been well identified in several heme-containing enzymes, such as coproheme decarboxylase (ChdC),^46^ heme-degrading enzymes from *Mycobacterium tuberculosis* (MhuD)^47,48^, and *S. aureus* (IsdI).^49^

### Thaxtomin B aromatic hydroxylation by TxtC

Following the TxtC-thaxtomin B MD simulation, three representative snapshots from MD trajectories were taken for the subsequent ONIOM calculations to investigate the aromatic thaxtomin B hydroxylation mechanism of TxtC. The most stable spin state for the corresponding reactant (RE2) remained in the doublet state. In ^2^RE2, as shown in Figure 4B, thaxtomin B placed its phenyl group over the heme ring with a dihedral angle between two molecular planes of 26.4°. The thaxtomin B C_20_ carbon atom was located at a distance of 3.51 Å from the heme γ-meso carbon. The steric interaction between thaxtomin B and heme caused ~1.20 Å of out-of-plane distortion on the heme plane (measured by normal coordinate structural decomposition (NSD)).^47,50^

Following Pathway-B, as shown in Scheme 1C, from ^2^RE2 the first thaxtomin B aromatic hydroxylation step was the addition of the phenyl group to the iron-oxo moiety, which proceeded on the doublet state potential energy surface by crossing over a transition state (^2^TS3) with a 15.4 kcal/mol energy barrier (see Figure 5A). By using the other two snapshots as the reactant initial structures, 16.7 kcal/mol and 16.8 kcal/mol energy barriers were obtained. The optimized ^2^TS3 geometric structure showed that the thaxtomin B phenyl group rotated by 4.5° around C_17_-C_18_ from that in ^2^RE2 so that the oxo moiety was located close to C_20_ within a distance of 1.97 Å. In ^2^TS3, C_20_ protruded from the phenyl plane to approach the oxyferryl moiety, while the Fe-O bond stretched to 1.73 Å. In ^2^TS3, the thaxtomin D phenyl group displayed a mixed cationic and radical character. At the ^2^TS3 stage, an electron from the substrate was partially transferred to the Cpd I moiety, as evidenced by the +0.31 charge carried by the thaxtomin B phenyl group. The electron transfer process from ^2^RE2 to ^2^TS3 could also be visualized from their electrostatic potential diagrams depicted in Figure S5.

**Figure 5.**
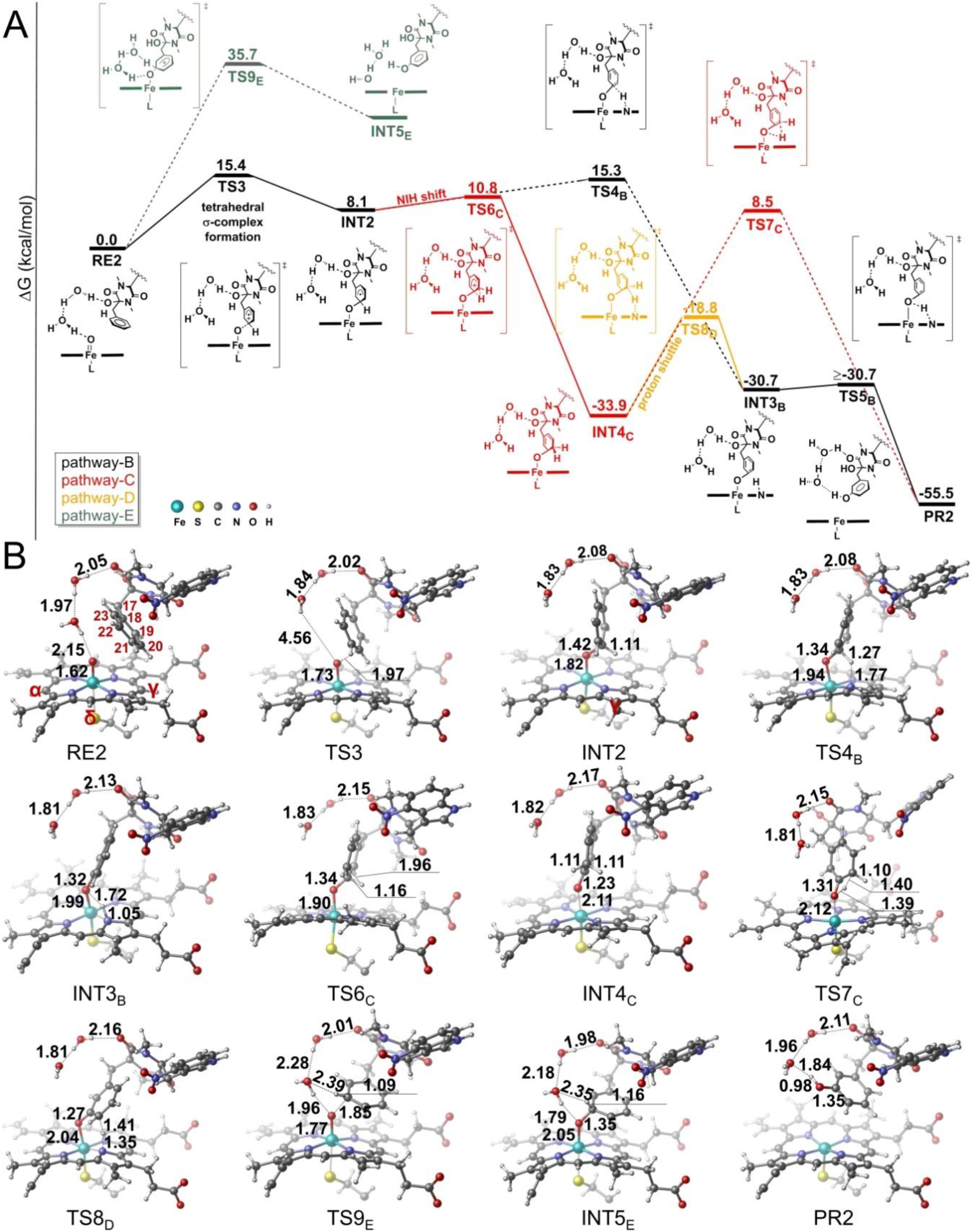
(A) Energy profile of the TxtC-catalyzed thaxtomin B aromatic hydroxylation process in doublet state. (B) Key structures of the stationary points along the aromatic hydroxylation in the TxtC doublet state. Bond lengths are given in Å.

**Table 2.** Calculated Mulliken spin population on various moieties in thaxtomin B aromatic hydroxylation. Por, Cys, thxB, Fe, O, N and H_C20_ represent the porphyrin, the distal Cys344 ligand, thaxtomin B, the iron center, Cpd I oxo moiety, porphyrin pyrrole nitrogen atom, and the hydrogen atom attached to C_20_, respectively.

| Compound | Por | Cys | thxB | Fe | O | N | H<sub>C20</sub> |
| --- | --- | --- | --- | --- | --- | --- | --- |
| <sup>2</sup>RE2 | -0.27 | -0.82 | 0.00 | 1.31 | 0.78 | -0.10 | 0.00 |
| <sup>2</sup>TS3 | -0.19 | -0.06 | -0.40 | 1.76 | -0.12 | -0.05 | -0.01 |
| <sup>2</sup>INT2 | -0.14 | 0.06 | -0.91 | 1.81 | 0.17 | -0.04 | -0.03 |
| <sup>2</sup>TS4<sub>B</sub> | -0.08 | 0.12 | 0.00 | 0.95 | 0.02 | -0.01 | 0.00 |
| <sup>2</sup>INT3<sub>B</sub> | -0.08 | 0.12 | 0.01 | 0.91 | 0.02 | -0.01 | 0.00 |
| <sup>2</sup>TS6<sub>C</sub> | -0.09 | 0.13 | -0.16 | 1.06 | 0.05 | -0.02 | 0.00 |
| <sup>2</sup>INT4<sub>C</sub> | -0.08 | 0.09 | 0.00 | 0.98 | 0.00 | -0.02 | 0.00 |
| <sup>2</sup>TS7<sub>C</sub> | -0.07 | 0.12 | 0.00 | 0.95 | 0.00 | -0.02 | 0.00 |
| <sup>2</sup>TS8<sub>D</sub> | -0.08 | 0.12 | 0.00 | 0.95 | 0.00 | -0.02 | 0.00 |
| <sup>2</sup>TS9<sub>E</sub> | -0.14 | -0.88 | 0.31 | 1.15 | 0.55 | -0.01 | 0.00 |
| <sup>2</sup>PR2 | -0.09 | 0.02 | 0.00 | 1.07 | 0.00 | -0.02 | 0.00 |

By crossing over ^2^TS3, the C_20_-O bond formed and generated the tetrahedral σ-complex intermediate ^2^INT2, which was endothermic at 8.1 kcal/mol. In σ-complex intermediate ^2^INT2, C_20_ converted to *sp^3^* hybridization with C_20_-H folding out of the phenyl plane towards the heme γ-meso carbon. The two C-C bonds of C_20_ with its adjacent carbon (C_20_-C_21_ and C_19_-C_20_) on the phenyl group both extended to 1.50 Å in ^2^INT2 (see Table S4). The next step was C_20_-H heterolysis, in which the proton was transferred to one of the porphyrin pyrrole nitrogen atoms to produce a N-protonated porphyrin complex (^2^INT3_B_), which went through transition state _2_TS4_B_ with a 7.2 kcal/mol local energy barrier (Figure 5A). The ^2^INT3_B_ geometric structure is shown in Figure 5B. Pyrrole nitrogen protonation caused severe porphyrin buckling so that the corresponding pyrrole fragment flipped out from the porphyrin plane. From ^2^INT3_B_, the proton on the pyrrole nitrogen could feasibly shuttle to the negatively charged oxygen atom of the nearby oxyferryl moiety, which resulted in full Fe-O bond cleavage and produced thaxtomin A (^2^PR2). This last transformation step took place on a flat area of the potential energy surface so that the reaction step was almost barrierless. It was similar to the benzene hydroxylation process catalyzed by cytochrome P450, which was reported in a previous study.^20^

From the tetrahedral σ-complex intermediate ^2^INT2, proton immigration from C_20_ to the O atom could proceed through an alternative pathway mediated by the phenyl group C_21_ para-carbon (NIH shift, pathway-C in Scheme 1C). This NIH shift process experienced an early stage transition state _2_TS6_C,_ which closely resembled ^2^INT2 with a low local barrier of 2.7 kcal/mol, in which the C_20_-H bond only stretched for 0.05 Å from that in ^2^INT2. After crossing proton immigration transition state ^2^TS6_C_, the resulting intermediate ^2^INT4_C_ energy dropped dramatically by 42.0 kcal/mol from _2_INT2. Such an exothermic process from ^2^INT2 to ^2^INT4_C_ could be rationalized by the fact that the substrate in ^2^INT2 phenyl group was in the form of a radical cation, as evidenced by a net - 0.91 spin population on it. This was quenched with the C_21_-H bond formation in ^2^INT4_C_, corresponding to a β electron transferred from the phenyl group to the oxyferryl moiety. In ^2^INT4_C_, the C_20_-C_21_ bond stretched to 1.51 Å, while C_20_-C_19_ was restored to 1.45 Å (see Table S4). From _2_INT4_C_, the proton shifting process could either proceed directly (pathway-C) or be mediated by the pyrrole nitrogen from the porphyrin ring (pathway-D). Our calculations showed that pathway-C needed to cross over a high energy barrier of 42.4 kcal/mol due to the severe steric tension of the rhombic shaped proton transfer transition structure (^2^TS7_C_, Figure 5). While pathway-D is the dominant one within which the reaction experiences the transition state ^2^TS8_D_ with energy barrier of 15.1 kcal/mol and generates ^2^INT3_B_ to merge in the pathway-B. It is worth noting that this three-stage pathway, which consisted of tetrahedral σ-complex intermediate (INT2) formation, NIH shift, and proton shuttle, bypassed the energetically unfavorable proton shift transition state ^2^TS4_B_ via protonation of the prophyrin pyrrole nitrogen, which therefore became the most favorable mechanism for thaxtomin B aromatic hydroxylation in TxtC.

### Heme distortion in TxtC

As discussed above, we explored the catalytic mechanisms of thaxtomin D aliphatic hydroxylation and thaxtomin B aromatic hydroxylation in TxtC. The calculations indicated that the thaxtomin B aromatic hydroxylation rate-limiting step experienced a slightly higher energy barrier than that of the thaxtomin D aliphatic hydroxylation (15.4 kcal/mol vs. 14.6 kcal/mol). However, a previous kinetic experiment determined that the TxtC aliphatic carbon hydroxylation catalytic efficiency was on the same order of aromatic hydroxylation with the corresponding energy barrier in the rate-limiting step of ~14.8 kcal/mol and ~15.0 kcal/mol, respectively.^3^ Thus, the calculations did not explain why TxtC displayed the observed regioselectivity to preferential hydroxylation on thaxtomin D aliphatic C_14_.

To elucidate the TxtC regioselectivity to preferentially hydroxylate thaxtomin D aliphatic C_14_, we further investigated the thaxtomin D aromatic hydroxylation process in TxtC with ONIOM calculations. The docked structure of complex A of TxtC-thaxtomin D after MD simulation was employed to build the initial reactant (RE2-D) and submitted for geometry optimization. An obvious structural characteristic of ^2^RE2-D, which was different from ^2^RE2 (the aromatic hydroxylation reactant of thaxtomin B in TxtC), was that the heme ring in it maintained a planar conformation. The measured NSD heme out-of-plane degree in RE2-D was only 0.34 Å. Based on RE2-D, we investigated the thaxtomin D aromatic hydroxylation process. We found that the calculated energy barrier of the rate-limiting step for generating the tetrahedral σ-complex intermediate (INT2-D) increased to 23.5 kcal/mol (By using the other two snapshots as the reactant initial structures, 24.9 kcal/mol and 29.4 kcal/mol energy barriers were obtained), which clearly suggested that without the hydroxyl group on C_14_, thaxtomin D aromatic hydroxylation would have been much slower. This calculation also rationalizes the observed TxtC regioselectivity to preferentially hydroxylate on the thaxtomin D aliphatic C_14_ site. A plausible explanation for the higher thaxtomin B reactivity on its aromatic site after hydroxylation on C_14_ could be ascribed to larger distortion of the heme in RE2 (NSD: 1.20 Å) than that in RE2-D (NSD: 0.34 Å). Our MD simulation revealed that the hydroxyl substituent on C_14_ induced several bulk water molecules to enter the active pocket, which pushed thaxtomin B towards the heme plane and caused considerable distortion. With that in mind, we manually removed the water molecules that were induced into the active pocket by the hydroxyl group on the C_14_ site and performed calculations on the rate-limiting tetrahedral σ-complex formation step on the TxtC-thaxtomin B complex again. As shown in Figure S6, without water molecules accumulating around C_14_ and pushing thaxtomin B to the heme plane, the heme ring was restored to a planar shape with the overall out-of-plane distortion degree descending to 0.66 Å after optimization. Correspondingly, the calculated energy barrier increased to 22.9 kcal/mol. Further computational studies were performed on a control model in which the water molecules around C_14_ were removed while the heme ring was restrained in a distortion conformation similar to that in RE2. The energy barrier of the rate-limiting step was calculated as 15.5 kcal/mol. Therefore, these calculations clearly revealed that the hydroxyl group on thaxtomin B C_14_ promoted its aromatic hydroxylation reactivity by causing distortion of the heme plane.

Our calculations revealed that thaxtomin B aromatic hydroxylation was rate-limited by the tetrahedral σ-complex formation step in which an α electron was transferred from the substrate phenyl group to Cpd I. To investigate why the heme out-of-plane distortion promoted the thaxtomin B aromatic hydroxylation reactivity in this step, it was intuitive to examine the reactant complex frontier orbitals (FMOs). Figure 6 depicts the lowest vacant RE2-D α-orbital. It presented as an anti-bonding molecular orbital that consisted of the a2u fragment orbital components from the porphyrin ring, the *dxz* orbital from the iron center, and the *px* orbital from the Cys344 ligand sulfur atom. The calculated energy level of this orbital was −4.43 eV, which was 1.02 eV higher than the highest occupied thaxtomin D α-orbital that populated the phenyl group. In ^2^RE2, the energy level of the lowest vacant α-orbital decreased to −4.86 eV. Although the highest occupied thaxtomin B α-orbital also decreased by 0.27 eV due to hydroxyl substitution on its C_14_ site, the energy gap still decreased to 0.86 eV. Obviously, such a narrower ^2^RE2 energy gap was consistent with its lower energy barrier to reach the tetrahedral σ-complex formation transition state. By analyzing the orbital components of their FMOs (Figure 6), one can find that the proportions of the fragment orbitals from the porphyrin, the iron center, and the sulfur atom of Cys344 ligand were changed from RE2-D to RE2 due to the heme out-of-plane distortion. In particular, the heme distortion decreased the anti-bonding overlap between the a2u orbital from porphyrin and *dxz* of the iron center, which therefore resulted in a lower energy level of this anti-bonding orbital. Similar orbital composition changes caused by heme out-of-plane distortion have also been observed in some heme-containing enzymes, such as ChdC, MhuD and IsdI. Based on the RE2 and RE2-D FMO analysis, we concluded that the heme out-of-plane distortion, which was caused by the hydrogen bond interaction between thaxtomin B and the surrounding water molecules induced by the C_14_ hydroxyl group, was crucial for the substrate aromatic hydroxylation by enhancing the Cpd I electrophilicity.

**Figure 6.**
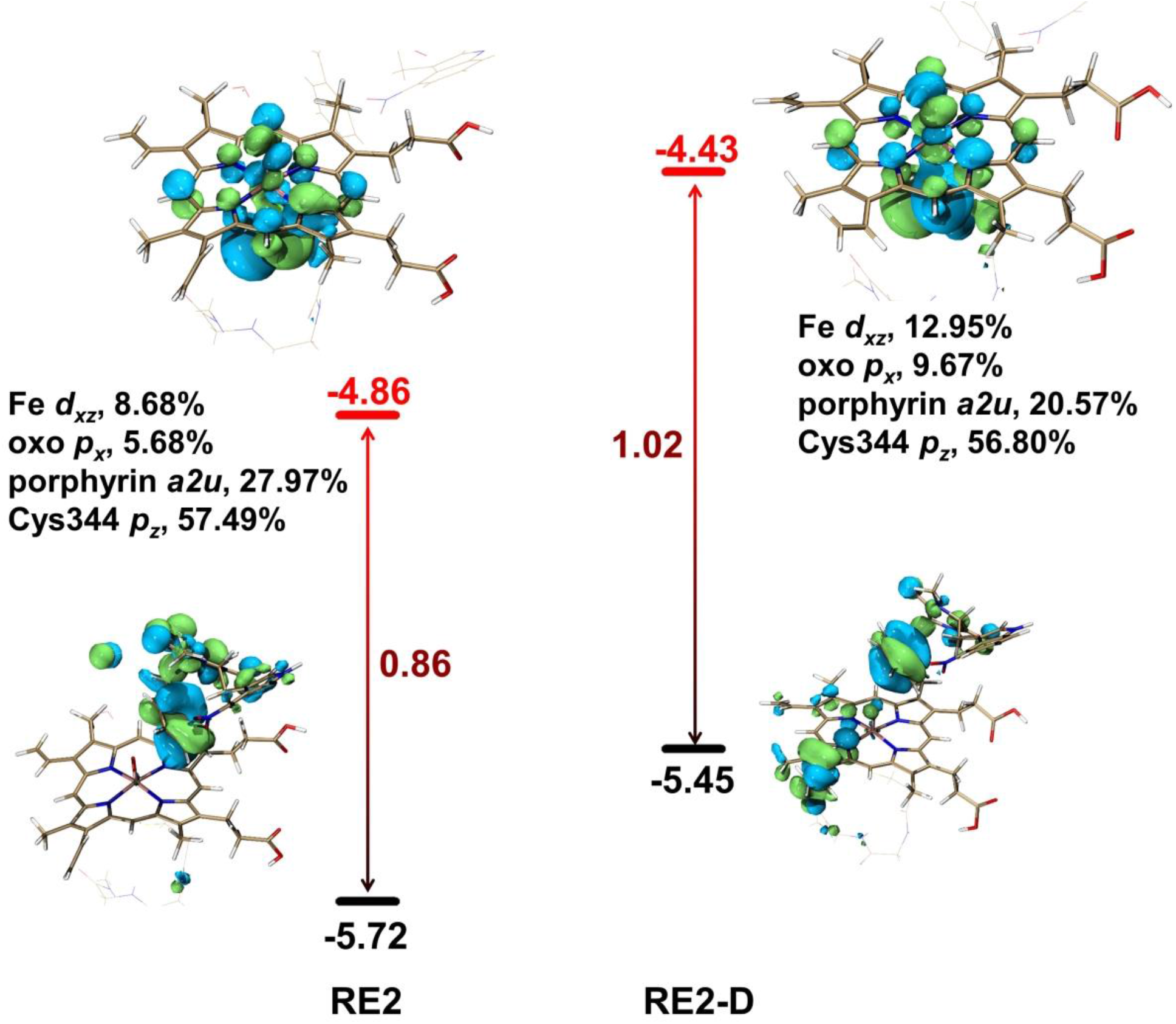
The highest occupied α-orbital of two substrates (black line) and the lowest Cpd I α-vacant orbital (red line) with their energies (in eV) for ^2^RE2 and ^2^RE2-D in the doublet state. The main compositions of the lowest Cpd I α-vacant orbital are also depicted (%).

## CONCLUSIONS

In this work, molecular docking and MD simulation showed that thaxtomin D was able to adopt a mixture of two conformations in the TxtC active site, in which either its aromatic C_20_-H or aliphatic C_14_-H lay towards the Cpd I active oxyferryl moiety. The thaxtomin D sequential aliphatic and aromatic hydroxylation mechanisms by TxtC were then investigated using detailed ONIOM calculations. The calculations revealed that the thaxtomin D C_14_ site aliphatic hydroxylation was 8.9 kcal/mol energetically favorable than its aromatic hydroxylation, which was consistent with the experimentally observed TxtC regioselectivity to hydroxylate on the thaxtomin D aliphatic C_14_ site. MD simulation on the complex of TxtC with thaxtomin B revealed that the hydroxyl group on the C_14_ site played a crucial role in promoting the substrate to flip over by placing the aromatic C_20_-H next to the oxyferryl center. The C_14_-OH group also induced the bulk water to enter the active site so that the substrate thaxtomin B was pushed towards the heme and resulted in the heme plane distortion, which finally accelerated the aromatic hydroxylation reaction by enhancing Cpd I electrophilicity.

## ACKNOWLEDGMENT

This work was supported by grants from the National Natural Science Foundation of China (no. 21571019, 21573020 and 21503018) and the National Key Research and Development Program of China (No. 2019YFC1805600).

## ABBREVIATIONS

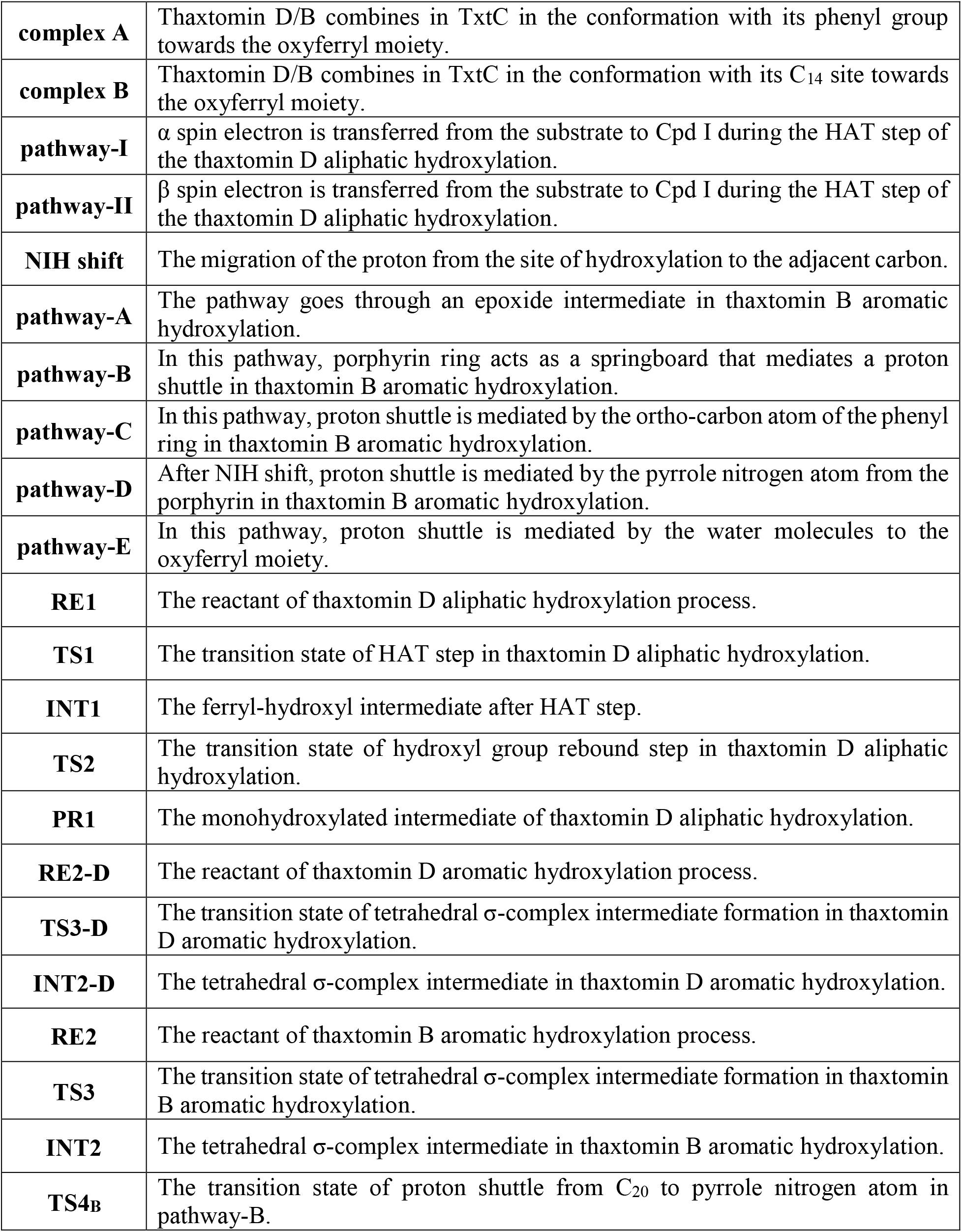
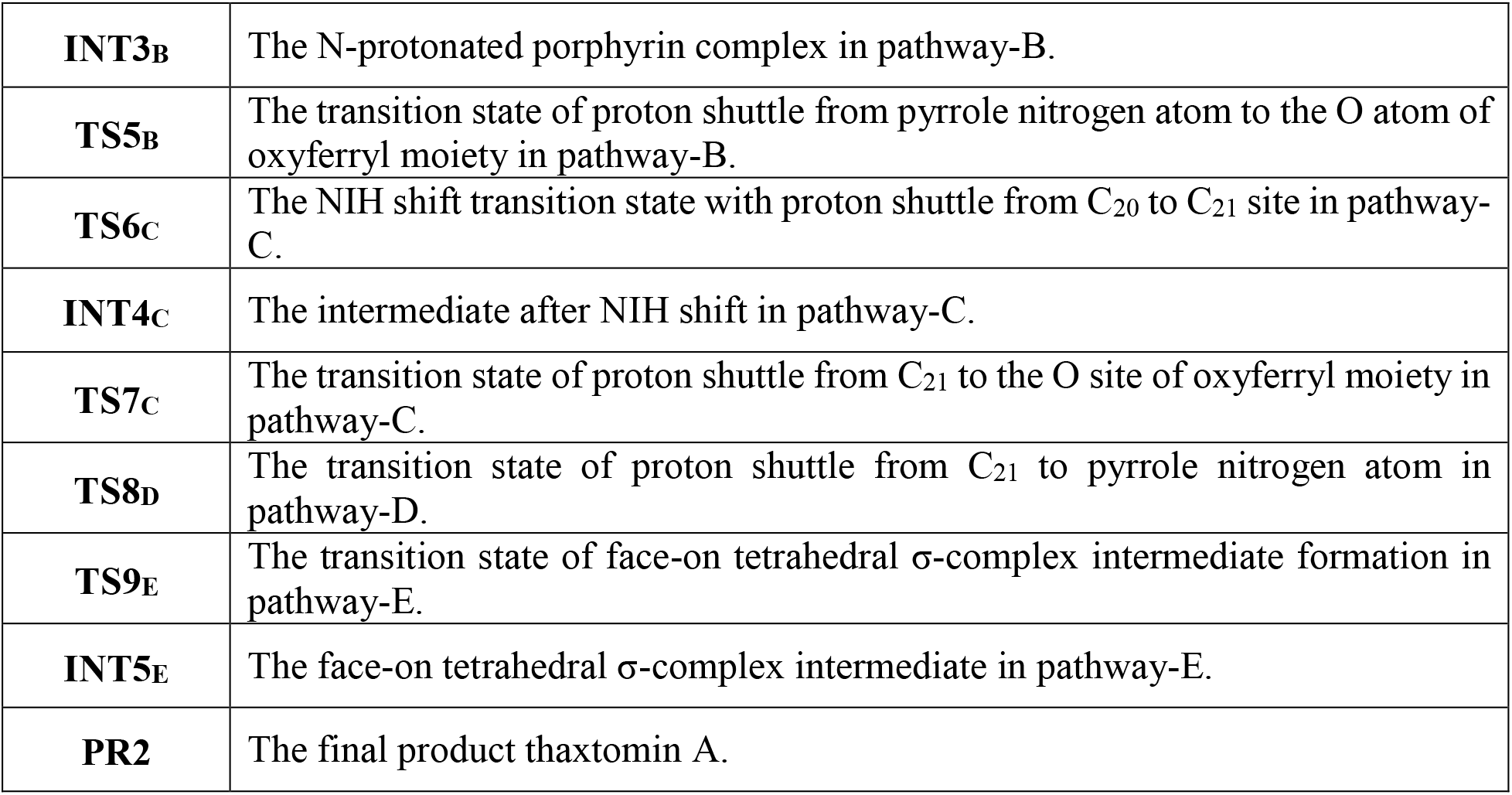

---

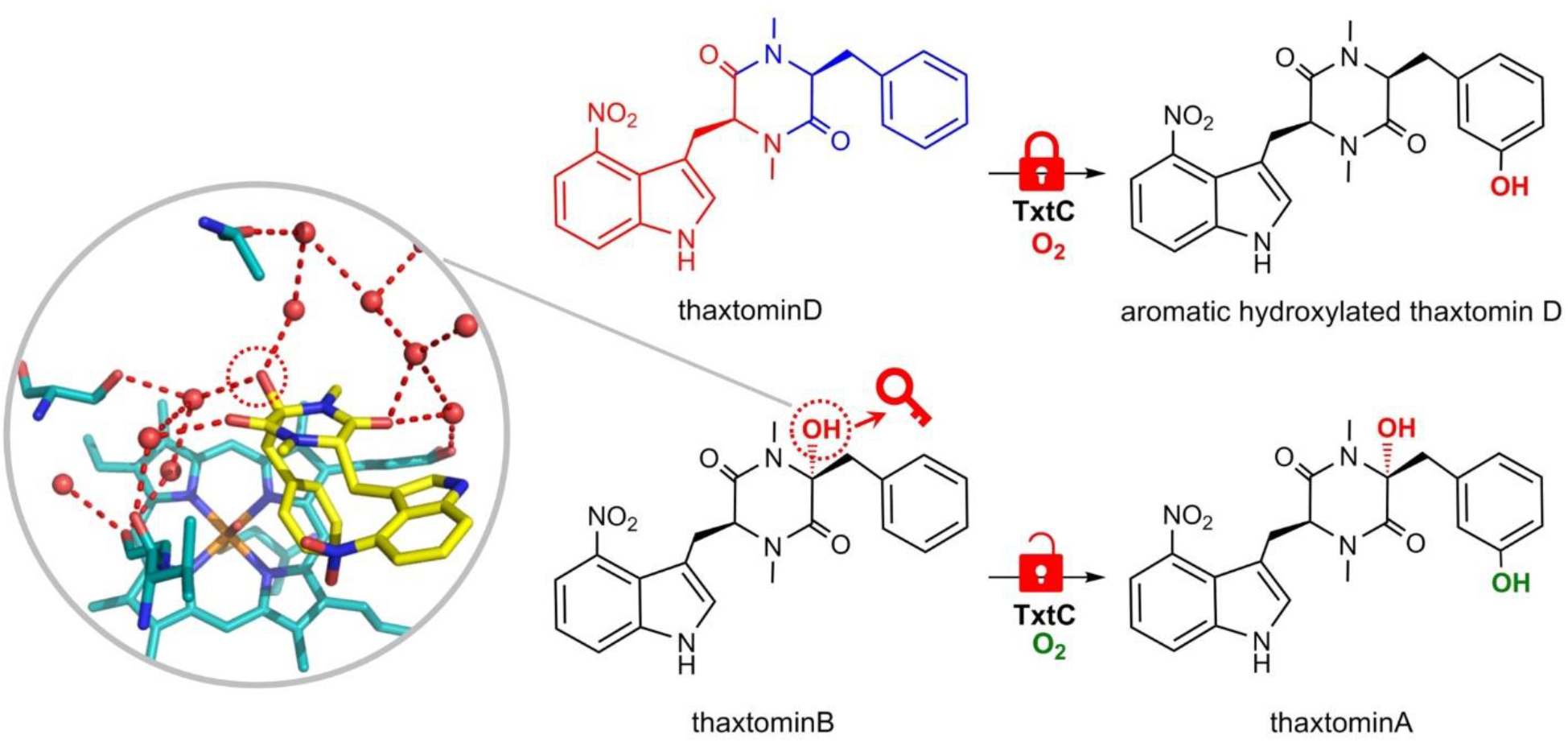

