## Supporting Information for "Interactive regulation between aliphatic hydroxylation and aromatic hydroxylation of thaxtomin D in TxtC: a theoretical investigation"

**Corresponding Authors:**

### 1 Details of the thaxtomin B aromatic hydroxylation in pathway-E

As indicated by the X-ray crystal structures of TxtC-thaxtomin D and TxtC-thaxtomin B, $C_{14}$ hydroxyl group induced two water molecules (WAT561 and WAT745) to enter into the active site and to form hydrogen bonds with Asp236, Thr385, as well as the substrate. Water molecules could play critical role in some enzymatic reactions. For example, water cluster network is responsible for the observed regioselectivity of heme hydroxylation in heme oxygenases.[1–3] In the N-demethylation process of N, N-dimethylanilines (DMAs) by cytochrome P450, water molecules also play the role to assist proton shift[4,5].

Considering that WAT745 is in close vicinity of the Fe-O site, we proposed an alternative thaxtomin B aromatic hydroxylation pathway with hydrogen on the $C_{20}$ shuttling through it to the Fe-O moiety (pathway-E, shown in Scheme 1D in the main text). In this pathway, the phenyl group of the substrate needs to rotate around $C_{17}$-$C_{18}$ bond for 20.5 degree via a face-on approach and lets the oxyferryl moiety attack from the other side of the phenyl plane so that the WAT745 is able to accept the proton delivered from $C_{20}$. Following this pathway, to generate the tetrahedral σ-complex intermediate ($^{2}$INT5$_{E}$), the reactant $^{2}$RE2 needs to cross over a 35.7 kcal/mol energy barrier, which in fact forbids the reaction to proceed via this pathway. Such a high energy barrier of this pathway is due to that the phenyl group of the substrate adopts a conformation being parallel to the heme plane, which results in less efficient orbital mixing between the Fe=O $\pi^*$ orbital of the oxyferryl moiety and the π orbital of the phenyl group moiety as indicated by Shaik in his work on the benzene hydroxylation by cytochrome P450[6].

### 2 Schemes

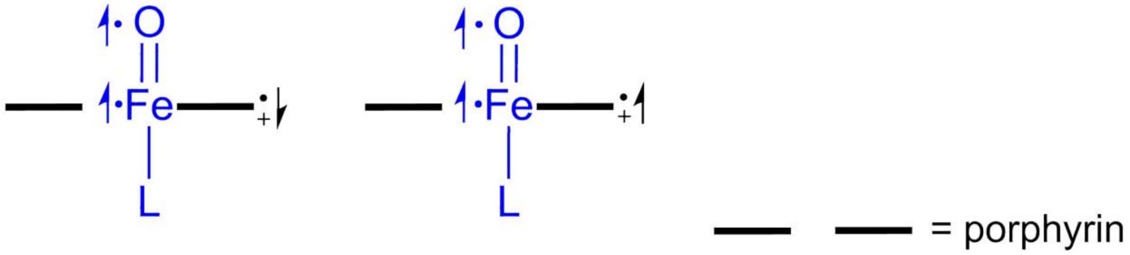


**Scheme S1.** The sketch map of doublet and quartet active oxyferryl species (Cpd I).

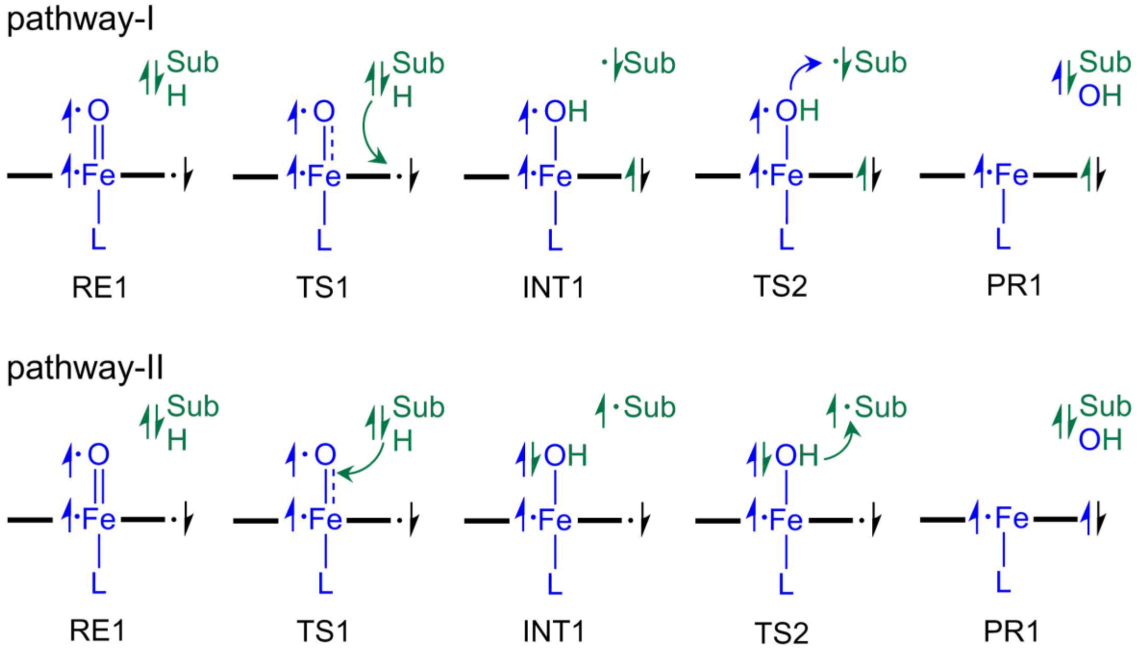


**Scheme S2.** Schematic representation of the two possible electron transfer pathways during the thaxtomin D aliphatic hydroxylation.

### 3 Figures

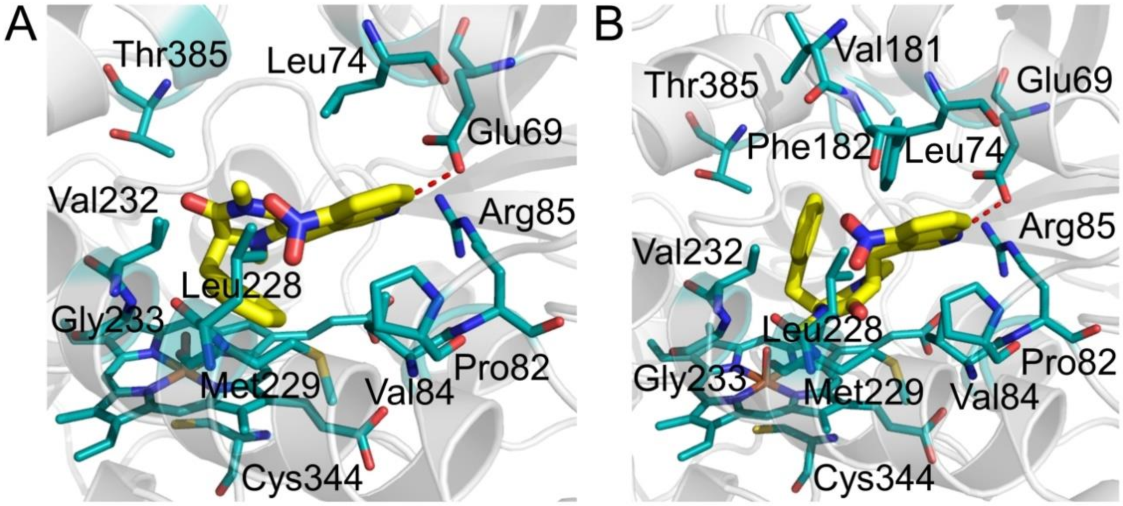


**Figure S1.** Two different conformations of thaxtomin D binding in the TxtC active site, complex A (left) and B (right), which were obtained from molecular docking. The key residues in the active site are represented as sticks. Carbon, oxygen, sulfur and nitrogen atoms are colored in cyan, red, orange and blue, respectively. The carbon atoms in the substrate molecules are colored in yellow. The hydrogen bonds are represented by red dashed lines.

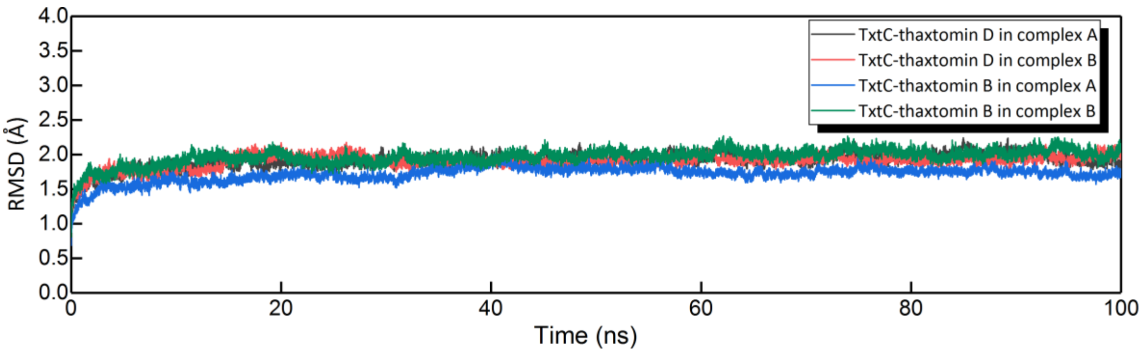


**Figure S2.** The root-mean-square deviation (RMSD) values for the TxtC-thaxtomin D and TxtC-thaxtomin B complexes during 100 ns simulation. In the complex A, thaxtomin D/B place the aliphatic $C_{14}$ next to the Fe-O moiety, while in the complex B, thaxtomin D/B place the aromatic $C_{20}$ next to the Fe-O moiety.

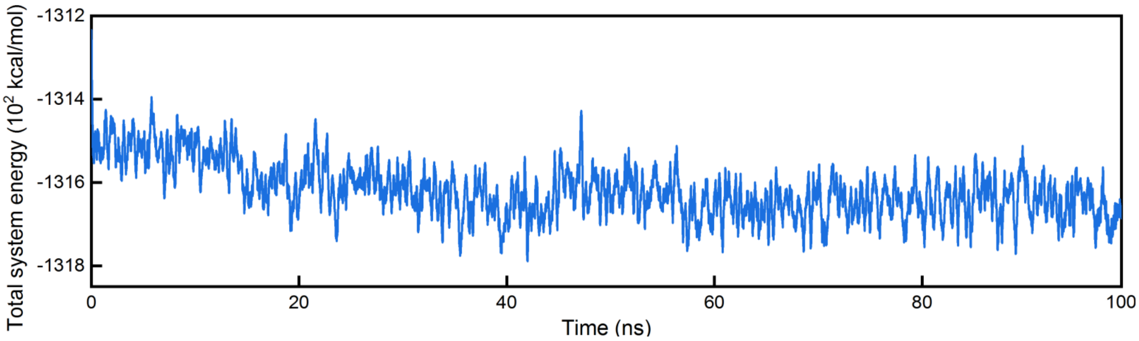


**Figure S3.** The moving average trend line of the total system energy during the 100 ns MD simulation.

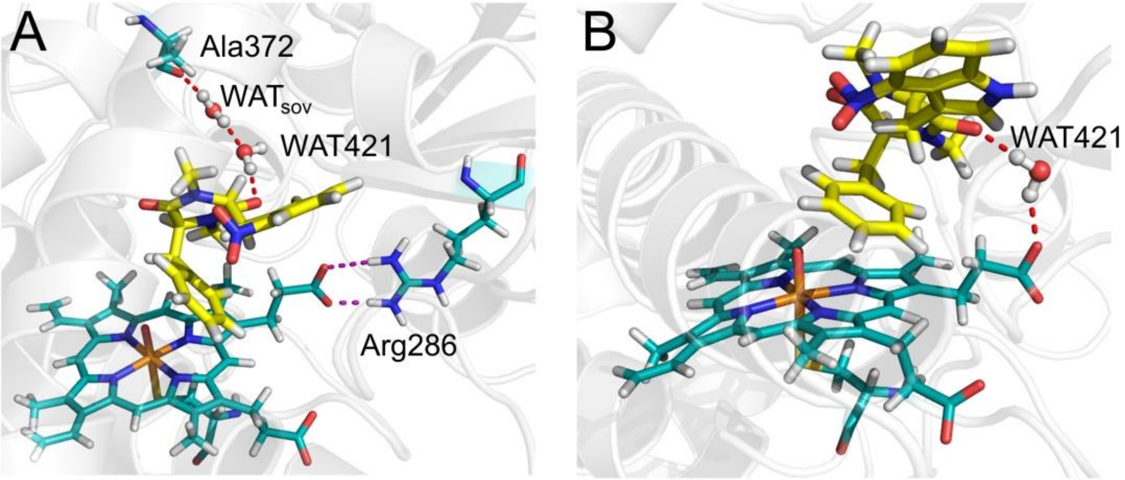


**Figure S4.** The snapshot of 100 ns from MD simulation (A) and the starting structure (B) of TxtC-thaxtomin D in the conformation of complex A. The key residues in the active site are represented as sticks. Carbon, oxygen, sulfur and nitrogen atoms are colored in cyan, red orange and blue, respectively. The carbon atoms in the substrate molecules are colored in yellow. The hydrogen bonds and salt bridge are represented by red and purple dashed lines, respectively.

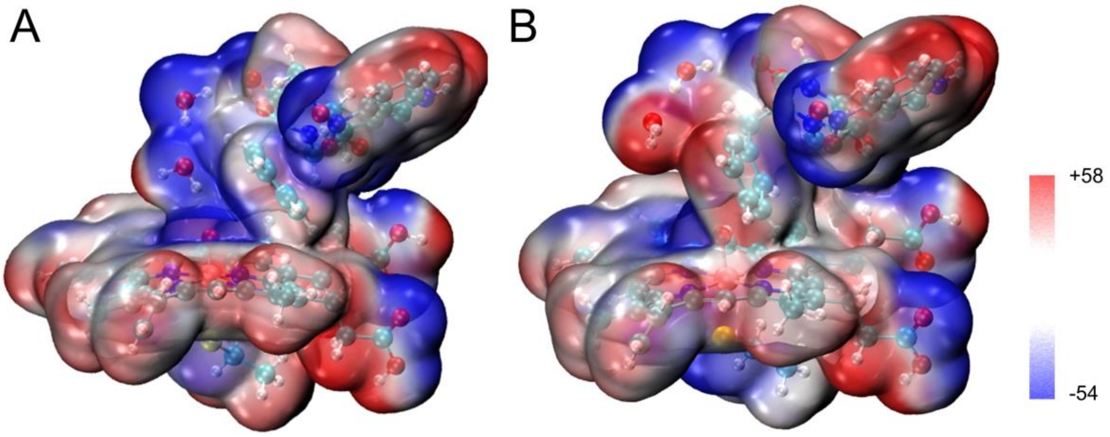


**Figure S5.** Electrostatic potential diagrams of the reactant ($^2$RE2) and the rate-limiting transition structure ($^2$TS3) in TxtC-catalyzed thaxtomin B aromatic hydroxylation, which clearly show that electronic charge is partially relocated from the thaxtomin B phenyl group to Cpd I.

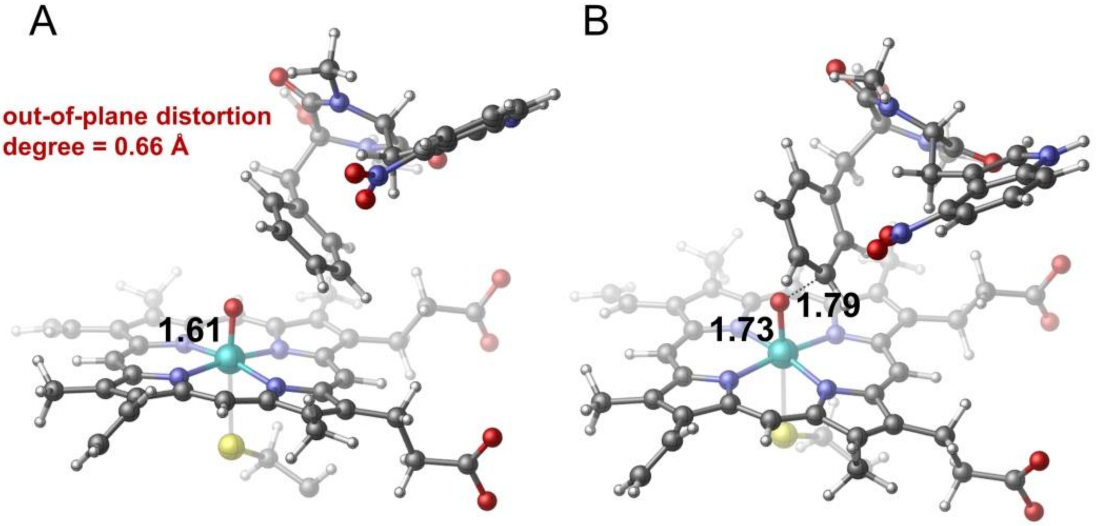


**Figure S6.** By removing the water molecules accumulated in the active site from the ONIOM model, the heme plane in the reactant (A) and rate-limiting transition state (B) tends to restore to a planar conformation.

### 4 Tables

**Table S1.** Calculated NPA charge densities on different moieties of the reaction species during aliphatic hydroxylation process of thaxtomin D. Fe, O, Cys, Por, thxD, $H_{C14}$ and $C_{14}$ represent the iron center, Cpd I oxo moiety, the distal Cys344 ligand, the porphyrin, thaxtomin D, the hydrogen atom attached to $C_{14}$, and the $C_{14}$ atom, respectively.

| Compound | Fe | O | Cys | thxD | Sub | $H_{C14}$ | $C_{14}$ |
|---|---|---|---|---|---|---|---|
| **RE1** | 0.32 | -0.32 | 0.09 | -0.09 | 0.00 | 0.27 | -0.13 |
| **TS1** | 0.41 | -0.59 | -0.06 | -0.17 | 0.41 | 0.40 | 0.04 |
| **INT1** | 0.44 | -0.71 | -0.02 | -0.20 | 0.49 | 0.48 | 0.21 |
| **TS2** | 0.84 | -0.22 | 0.05 | -0.15 | -0.01 | 0.24 | -0.06 |
| **PR1** | 0.38 | -0.71 | 0.03 | -0.42 | 0.72 | 0.50 | 0.35 |

**Table S2.** Calculated NPA charge densities on different moieties of the reaction species during aromatic hydroxylation process of thaxtomin B. Por, Cys, thxB, Fe, O, N and $H_{C20}$ represent the porphyrin, the distal Cys344 ligand, thaxtomin B, the iron center, Cpd I oxo moiety, porphyrin pyrrole nitrogen atom, and the hydrogen atom attached to $C_{20}$, respectively.

| Compound | Por | Cys | thxB | Fe | O | N | $H_{C20}$ |
|---|---|---|---|---|---|---|---|
| **RE2** | -0.10 | 0.12 | -0.01 | 0.29 | -0.32 | -0.36 | 0.20 |
| **TS3** | -0.15 | -0.01 | 0.31 | 0.24 | -0.40 | -0.36 | 0.24 |
| **INT2** | -0.15 | 0.10 | 0.25 | 0.27 | -0.48 | -0.37 | 0.20 |
| **$TS4_B$** | -0.31 | -0.02 | 0.75 | 0.15 | -0.59 | -0.43 | 0.38 |
| **$INT5_B$** | -0.11 | 0.02 | 0.57 | 0.16 | -0.65 | -0.54 | 0.49 |
| **$TS6_C$** | -0.42 | -0.01 | 0.81 | 0.18 | -0.58 | -0.37 | 0.30 |
| **$INT6_C$** | -0.43 | 0.08 | 0.67 | 0.15 | -0.49 | -0.37 | 0.27 |
| **$TS7_C$** | -0.25 | -0.05 | 0.36 | 0.73 | -0.30 | -0.20 | 0.23 |
| **$TS8_D$** | -0.24 | 0.04 | 0.60 | 0.18 | -0.60 | -0.51 | 0.41 |
| **$TS9_E$** | -0.28 | 0.20 | 0.24 | 0.35 | -0.51 | -0.38 | 0.22 |
| **PR2** | -0.37 | 0.07 | 0.63 | 0.30 | -0.67 | -0.39 | 0.21 |

**Table S3.** Relative energies (in kcal/mol) of the reaction intermediates and transition states on quartet spin-state surface for the thaxtomin B aromatic hydroxylation.

| **RE2** | **TS3** | **INT2** | **TS4$_B$** | **INT3$_B$** | **TS6$_C$** |
|---|---|---|---|---|---|
| 1.3 | 24.6 | 10.1 | 22.4 | -29.2 | 23.2 |
| **INT4$_C$** | **TS7$_C$** | **TS8$_D$** | **TS9$_E$** | **PR2** | **--** |
| -33.2 | 19.5 | -12.33 | 27.14 | -56.3 | **--** |

**Table S4.** The C-C bond lengths (unit for distances is angstrom) of thaxtomin B phenyl group in optimized reaction species along its aromatic hydroxylation process. Atom indexes refer to Figure 4B in the main text.

| **Compound** | **$C_{18}$-$C_{19}$** | **$C_{19}$-$C_{20}$** | **$C_{20}$-$C_{21}$** | **$C_{21}$-$C_{22}$** | **$C_{22}$-$C_{23}$** | **$C_{23}$-$C_{18}$** |
|---|---|---|---|---|---|---|
| **RE2** | 1.40 | 1.40 | 1.39 | 1.40 | 1.40 | 1.40 |
| **TS3** | 1.39 | 1.42 | 1.42 | 1.38 | 1.40 | 1.42 |
| **INT2** | 1.37 | 1.50 | 1.50 | 1.37 | 1.42 | 1.43 |
| **TS4$_B$** | 1.38 | 1.46 | 1.45 | 1.39 | 1.40 | 1.42 |
| **INT3$_B$** | 1.40 | 1.42 | 1.42 | 1.39 | 1.40 | 1.41 |
| **TS6$_C$** | 1.38 | 1.48 | 1.48 | 1.37 | 1.41 | 1.42 |
| **INT4$_C$** | 1.37 | 1.45 | 1.51 | 1.49 | 1.35 | 1.47 |
| **TS7$_C$** | 1.40 | 1.40 | 1.44 | 1.43 | 1.38 | 1.44 |
| **TS8$_D$** | 1.38 | 1.43 | 1.45 | 1.44 | 1.38 | 1.43 |
| **TS9$_E$** | 1.42 | 1.41 | 1.38 | 1.43 | 1.44 | 1.39 |
| **PR2** | 1.40 | 1.40 | 1.39 | 1.40 | 1.40 | 1.40 |
